# Predicting the Activities of Drug Excipients on Biological Targets using One-Shot Learning

**DOI:** 10.1101/2021.10.04.463110

**Authors:** Xuenan Mi, Diwakar Shukla

**Affiliations:** Center for Biophysics and Quantitative Biology, University of Illinois at Urbana-Champaign,Urbana, IL 61801, USA; Department of Chemical and Biomolecular Engineering, University of Illinois at Urbana-Champaign,Urbana, IL 61801, USA; Beckman Institute for Advanced Science and Technology, University of Illinois at Urbana-Champaign,Urbana, IL 61801, USA; Cancer Center at Illinois, University of Illinois at Urbana-Champaign,Urbana, IL 61801, USA; Center for Digital Agriculture, University of Illinois at Urbana-Champaign,Urbana, IL 61801, USA; Department of Plant Biology, University of Illinois at Urbana-Champaign,Urbana, IL 61801, USA; National Center for Supercomputing Applications, University of Illinois at Urbana-Champaign,Urbana, IL 61801, USA

## Abstract

Excipients are a major component of drugs and are used to improve drugs attributes such as stability and appearance. Excipients approved by Food and Drug Administration (FDA) are regarded as safe for human in allowed concentration, but their potential interaction with drug targets have not been investigated systematically, which might influence drug’s efficacy. Deep learning models have been used for identification of ligands that could bind to the drug targets. However, due to the limited available data, it is challenging to reliably estimate the likelihood of a ligand-protein interaction. One-shot learning techniques provide a potential approach to address this low-data problem as these techniques require only one or a few examples to classify the new data. In this study, we apply one-shot learning models on datasets that include ligands binding to G-Protein Coupled Receptors (GPCRs) and Kinases. The predicted results suggest that one-shot learning models could be used for predicting ligand-protein interaction and the models attain better performance when protein targets contain conserved binding pockets. The trained models are also used to predict interactions between excipients and drug targets, which provides a potential efficient strategy to explore the activities of drug excipients. We find that a large number of drug excipients could interact with biological targets and influence their function. The results demonstrate how one-shot learning models can be used to make accurate prediction for excipient-protein interactions and these methods could be used for selecting excipients with limited drug-protein interactions.

## Introduction

Drug excipients, also called inactive ingredients, are compounds approved by the US Food and Drug Administration (FDA), which play important non-pharmaceutical effects in formulation development. The active pharmaceutical ingredients (API) are components in drugs that provide the pharmaceutical effect, but excipients are used for improving physical properties of drugs essential for efficient delivery, stability and bioavailability. Some examples of drug excipients include sucrose and trehalose, which function as conformational stabilizer; Arginine hydrochloride, which is used as an aggregation suppressor; D&C Red No. 28, FD&C Yellow No.5, are used as dyes for coloring drugs so that they can be distinguished easily; Butylated hydroxytoluene are antioxidant for improving drug’s shelf life.^1^ Although approved excipients do not exhibit apparent toxicity in animal studies and clinical use, they are likely to interact with other biological targets, which may influence the drug’s therapeutic effect or even result in unwanted side effects. Recent studies have reported excipients in oral medications could cause adverse reactions.^2^ A total of 38 inactive ingredients such as chemical dyes and lactose have been found to cause potential allergic symptoms. The systematic identification of potential targets of excipients is necessary to avoid the unwanted side effects caused by excipients added in the drugs. Therefore, as a major component of drug mass, the effect of excipients on target proteins needs to be investigated for efficient drug formulation development. Detailed computational and experimental investigations of excipients have been performed for assessing the relationship between chemical structure of excipients and their performance^3–6^ but their impact on other proteins in the body have not been systematically investigated or even considered during the formulation development process. The computational cost of investigating the impact of the complex mixture of chemicals on different proteins is both experimentally and computationally challenging^7–9^. Therefore, there is a need to develop methods which could enable rapid and reliable prediction of the excipient-protein interactions

Machine Learning (ML) methods have been used to predict the biological effect of excipients and Generally recognized as safe (GRAS) compounds. Reker et al. have used random forest models to reveal the unknown biological effects of inactive ingredients.^10^ While traditional ML approaches have provided useful insights in to biological activity of excipients, deep learning methods have the potential provide a more accurate estimate of excipient-protein interactions. Deep learning is a subclass of machine learning algorithms which uses multi-layer neural network architectures to learn representations of data. This architecture has dramatically improved the performance of ML methods in speech recognition, computer vision, and object detection.^11^ In the last decade, machine learning models have been increasingly applied for drug discovery. In 2012, Merck organized the Molecular Activity Challenge, aimed to identify the best model for predicting biological activities of different molecules based on their chemical structures. The challenge was won by a multitask deep network that increases relative prediction accuracy by 15% over the baseline.^12^ In addition to molecular activity prediction, deep learning has also remarkable performance in drug-target interaction prediction,^13,14^ molecular de novo design,^15^ and chemical syntheses.^16^ However, the lack of large labeled datasets for drug design has limited the effectiveness of these innovated deep neural methods in drug discovery. Typically, the training of deep learning models with large number of layers requires large amounts of data. For example, one of the most popular deep learning datasets, ImageNet, contains more than 14 million images.^17^ Therefore, there is a need to develop and employ methods that could use small and sparse datasets in the drug discovery projects. Several recent works have integrated multiple data sources and utilized the transferability between them to address this low data issue.^18,19^

One of the solutions to low data problem is one-shot learning, which classifies new data having seen only one or a few training examples. The term one-shot learning was proposed by Fei-Fei Li in 2006,^20^ which takes advantage of the knowledge from previously learned categories, even though these categories are different from the target category. Recent advances in one-shot learning is in combination with deep learning, which learns a meaningful distance metric through comparing new data point to the limited inputs. The metric-based one-shot learning was first implemented in the Siamese Neural Network^21^ designed for image recognition. A remarkable improvement of one-shot learning was made by Matching Network^22^ approach, in which the feature embedding was improved such that the architecture could extract prior knowledge better. In 2017, based on the Matching Network, researchers developed the Iterative Refinement Long Short-Term memory (IterRefLSTM) approach that was recently adapted to the drug discovery purpose.^23^

In this work, we apply one-shot learning networks for predicting drug excipients binding to G protein-coupled receptors (GPCRs) and protein kinases. GPCRs are a family of integral membrane proteins that play crucial role in diverse cellular and biological activities. GPCRs are the most intensively studied drug targets and account for around 27% of the global pharmaceutical market.^24^ Protein kinases are the second class of proteins targeted for drug discovery. Kinases are enzymes that transfer phosphate group to a protein and are associated with human cancer, immunological and degenerative disease.^25^ We apply one-shot learning models on GPCR and Kinase ligand datasets, and find the model with high prediction performance even with limited data. Finally, the trained model was used for predicting the excipients binding to the GPCRs and protein kinases. The predicted results may accelerate the discovery or design noval of drug excipients and help shift the paradigm of pharmaceutical formulation development from experiment-dependent studies to data-driven methodologies.

## Methods

### Datasets Curation

We developed one-shot learning models using two datasets: the kinase-inhibitor and the GPCR-ligand datasets. The kinase-inhibitor dataset include 420 unique kinase targets, 36, 628 inhibitors and 123, 005 kinase-inhibtor interactions.^26^ The GPCR-ligand dataset was downloaded from GPCRdb (https://gpcrdb.org). As the largest sub-family of GPCR, 525 class A GPCRs, 132, 354 ligands and 215, 684 GPCR-ligand bindings were retrieved from GPCRdb. We use a simplified molecular-input line-entry system (SMILES) of each ligand and inhibitor in which a unique SMILES string is mapped to a compound structure.^27^ All data collected are treated as positive samples for training and negative samples are not defined in the datasets. We generate the negative samples following the steps below:

Step 1. Generate all receptor pairs (receptor A, receptor B) which have no overlapping ligands

Step 2. In each pair, swap all ligands of receptor A with ligands of receptor B, and make negative samples using ligands of receptor B with receptor A **OR** ligands of receptor A with receptor B

Step 3. Randomly select negative samples from the pool generated from step 2

### One-shot Learning Architectures

In this study, we considered kinase-inhibitor and GPCR-ligand data as a collection of multiple binary learning tasks. For example, in the kinase dataset, each kinase is treated as one binary task, and in each task known binding and non-binding compounds are treated as positive examples and negative examples, respectively. Some tasks in the dataset are used for training a one-shot learning model. The goal is to create a strong classifier for the remaining tasks in the testing set. In other words, we train the model using the data from the subset of kinases and GPCRs to predict the ligand activity for other kinases and GPCRs respectively. The following sections describe the three one-shot learning models used in this study: Siamese network,^21^ Matching network,^22^ and Iterative Refinement LSTM.^23^ Figure 1 shows the schematic architecture of these one-shot learning techniques. More details on each architecture are provided in the following sections.

**Figure 1:**
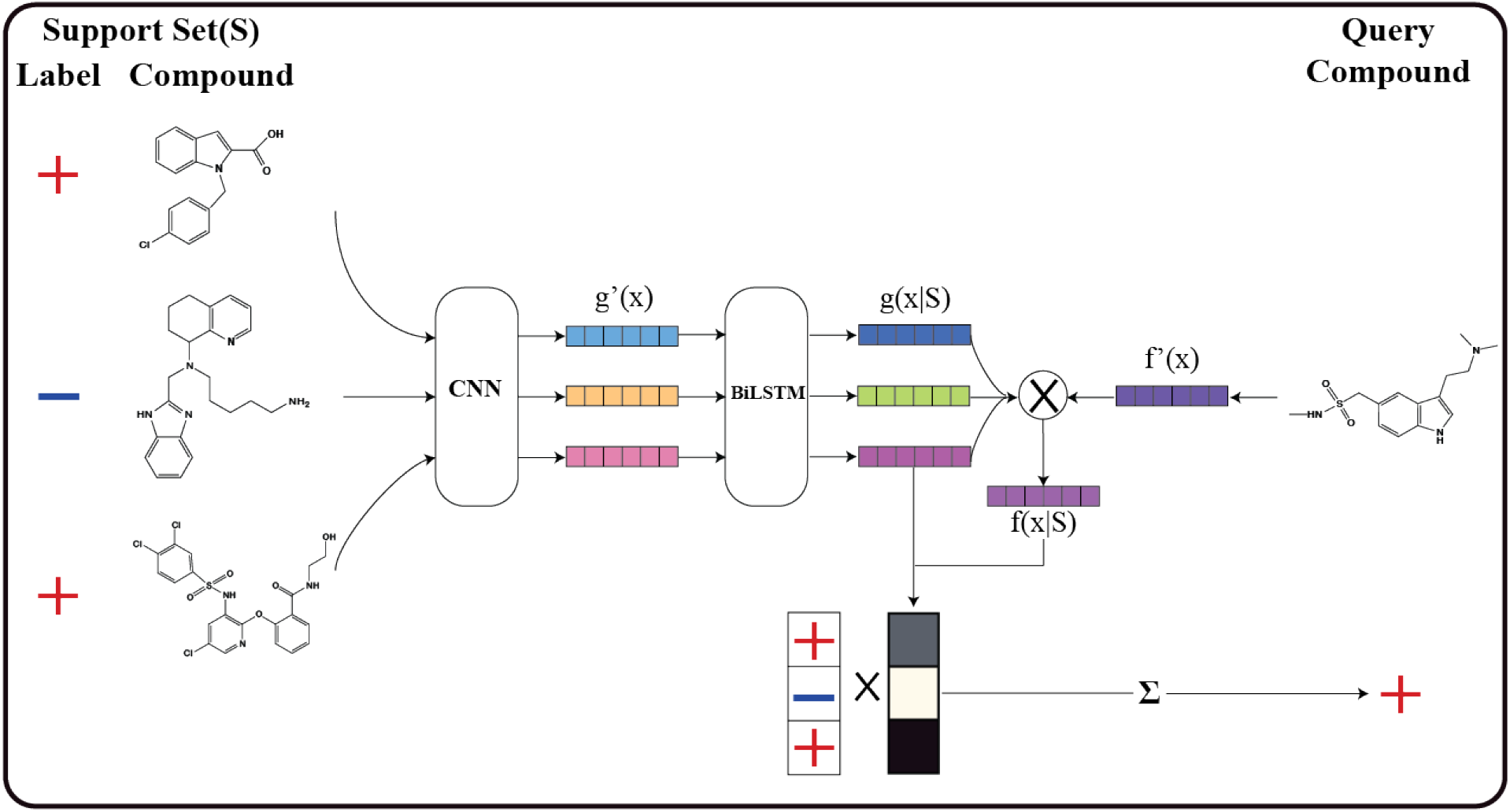
The schematic architecture of one-shot learning model. For the structure of each compound in the support set *S*, the initial embedding *g ′* (*x*) is generated from standard convolutional neural network (CNN). Then the initial embedding is passed through bidirectional LSTM (BiLSTM), generating full context embedding *g*(*x* | *S*) for each support data. For query structure, the full context embedding *f* (*x* | *S*) is produced from the initial embedding for query *f* ′ (*x*) and *g*(*x* | *S*) based on the attention mechanism. The similarity between *f* (*x* | *S*) and each *g*(*x* | *S*) is calculated, and the predicted label for query structure is generated based on the similarity distribution over *S*.

### Siamese Network

The Siamese network^21^ is the simplest one-shot model, which computes the distance between query data and each data point in the support set and then performs a weighted combination of support set labels. In the Siamese network, the two inputs (query data and each element in support set) are passed through the same convolutional neural network with shared weights at each hidden layer and generate a fixed length feature vector. The distance between two feature vectors are calculated and can be interpreted as dissimilarity between the query compound and the compound in the support set. Based on the generated similarity for each compound in support set, the different weight will be assigned to corresponding support set label. The predicted label for query compound will be generated through combining the weighted support set labels.

Mathematically, the predicted label ŷ for query compound *x* is defined as Equation (1), the function *a*(*x, x*_*i*_) denotes the weighting function for query compound *x* and support set element *x*_*i*_ with associated label *y*_*i*_. The weight function *a* can be interpreted as distribution over the support set where the sum of weights 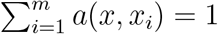.

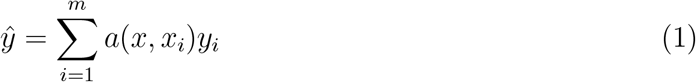

The weight function *a*(*x, x*_*i*_) can be computed as Equation (2), *f* (*x*) denotes embedding function of the query compound, while *g*(*x*_*i*_) denotes embedding function of a compound in support set. Here, *f* and *g* are graph convolutional networks and in the Siamese network, *f* and *g* are the identical function. The similarity between two vectors are represented by *k*(*f* (*x*), *g*(*x*_*j*_))

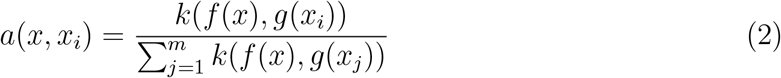

### Matching network (Attention LSTM)

Compared to the Siamese network, an important contribution of matching network^22^ is using full context embedding *g*(*x*_*i*_|*S*) and *f* (*x*|*S*) instead of independent embedding *g*(*x*_*i*_) and *f* (*x*). In the simple one-shot model, Siamese Network, the embedding of query compound *f* (*x*) and each support compound *g*(*x*_*i*_) is computed independently of the whole support set *S*. The effect of the context of the entire support set on the embeddings of query and each individual support compound is ignored. The full context embeddings address this challenge, which allows the embedding for *x* and *x*_*i*_ to influence each other.

In order to obtain the full context embedding *g*(*x*_*i*_|*S*), the matching network utilizes the bidirectional Long-Short Term Memory (BiLSTM)^28^ to encode *x*_*i*_ given the all information of support set *S*. The BiLSTM architecture allows each *g*(*x*_*i*_) will be affected by all elements in the support set. *g*(*x*_*i*_|*S*) is define as follows:

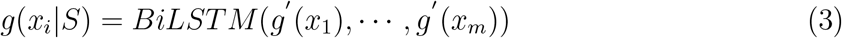

where *g ′* is the initial embedding for element in support set using graph convolutional neural network. The LSTM network is a kind of recurrent neural network, which is used for processing sequential input. The BiLSTM is used, instead of LSTM, to partially reduce the dependence on the order of input, which will improve the performance of one-shot models because there is no ordering of elements in the support set. However, BiLSTM cannot remove all dependence on order of input.

For query compound *x*, the matching network defines a full embedding function based on attention LSTM model (attLSTM):^29^

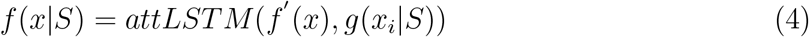

where *f ′* is the initial embedding for query compound using the graph convolutional neural network which are input to the LSTM. In the attLSTM, there are K times of unrolling steps of LSTM, which allows for transfer information among inputs. This algorithm will enhance the influence of important element in the support set on the query data and ignore unrelated elements. The attLSTM is an order-independent mechanism. As there is no ordering of elements in support set, order-independent mechanism is more reasonable for our problem.

### Iteration Refinement LSTM Network

BiLSTM only partially removes the order-dependence in full context embedding *g*(*x*_*i*_|*S*) for each element in the support set. As there is no natural order in support set, so any order-dependence is not ideal. In addition, the full context embedding *g*(*x*_*i*_|*S*) for support compound and *f* (*x*|*S*) for query is not symmetric. *g*(*x*_*i*_|*S*) depends only on all *g ′* (*x*), but *f* (*x*|*S*) depends on *f ′* (*x*) and *g*(*·*|*S*). This kind of asymmetry indicates some additional information is ignored, thus will influence the model performance.

In 2017, a new architecture called iterative refinement LSTM was proposed.^23^ In this architecture, attLSTM is used to generate both *g*(*x*_*i*_|*S*) and *f* (*x*|*S*) which solves the order dependence issue in *g*(*x*_*i*_|*S*) and asymmetric property of support and query embedding. First, the initial embeddings *f ′* (*x*) and *g ′* (*x*) are generated from the graph convolutional neural network. Then briefly, the query embedding *f* and support embedding *g* are updated iteratively based on the attention mechanism and allows the embedding of the support set to influence the query embedding iteratively.

### Model Training and Evaluation

Let us define a Task *T* as one of all learning tasks. We split *T* into two sets, Train-Tasks and Test-tasks. Let S represent a support set, let B represent a batch of queries. Training consists of a sequence of episodes. In each episode, a task *T* is randomly sampled, and then a support *S* and a batch of queries *B* are sampled from the task. The models is trained to minimize the error predicting the labeles in the batch *B* conditioned on the support set *S*, as described in Equation (5):

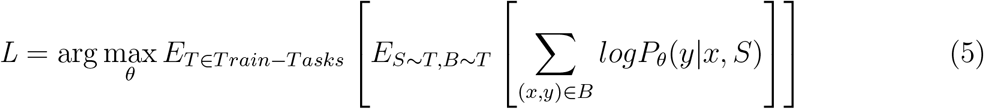

Model performance was evaluated for each task in the testing set. For each test task, a support set is sampled randomly from data points in that task and the AUC score for the model was evaluated on the remaining data points (excluding the support data) in that task. This procedure was repeated 20 times and the mean and standard deviation of AUC score of each test task was calculated.

## Results

### One Shot Learning Methods Provide Accurate Prediction of Kinase-Ligand Interaction

The Kinase dataste consists of 420 kinase with 36, 628 inhibitors, forming 123, 005 unique kinase-inhibitor interactions. We observe that majority of protein kinase have a small number of known inhibitors (Figure 2a), among them, 50% of protein kinase have less than 50 known inhibitors and 16% kinase have less than 10 inhibitors. The majority of kinases have few experimentally verified inhibitors, so the traditional deep learning networks are limited by the current data availability for prediction of ligand binding.

**Figure 2:**
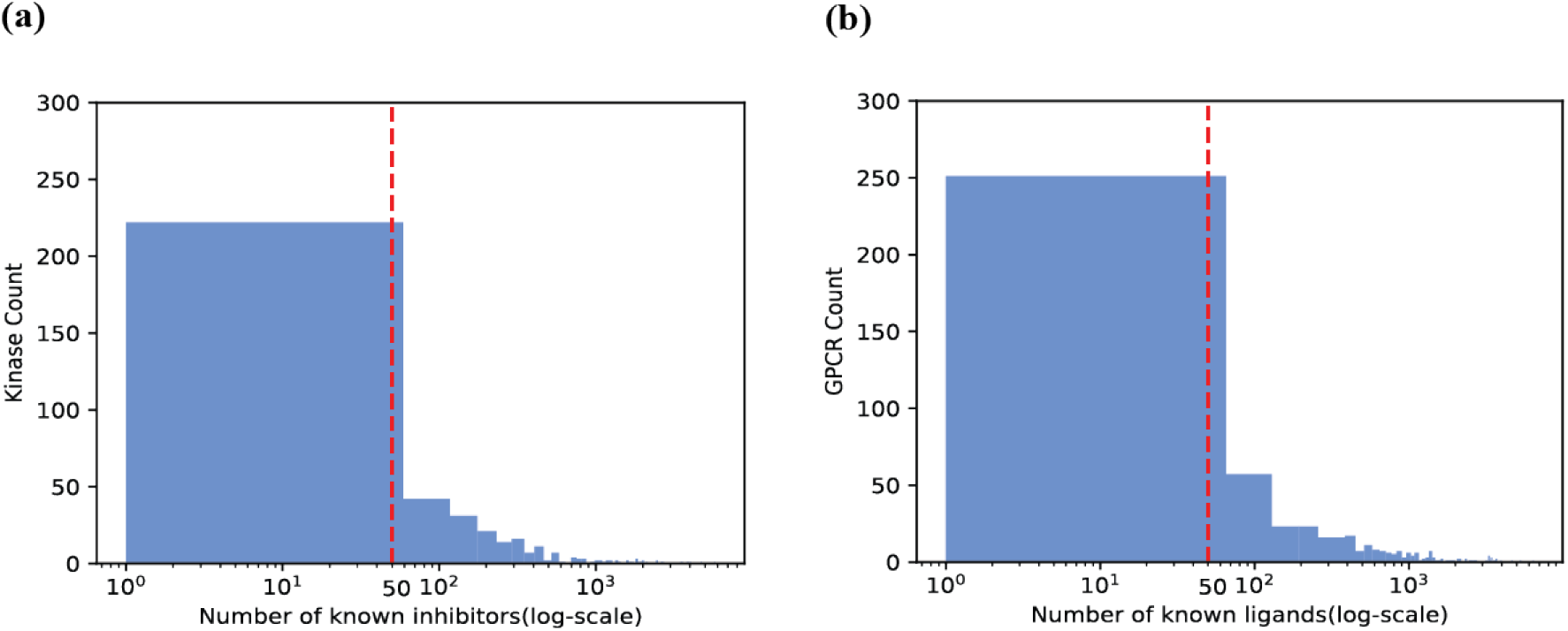
A summary of data distribution of Kinase-inhibitor collection and GPCR-ligand collection. (*a*) A majority protein kinase have a small amount of known inhibitors. The left side of red dashed line represents the protein kinase with *<* 50 known inhibitors, which accounts for 50.23% of total protein kinase. (*b*) Most GPCRs have limited known binding ligands. The left side of red dashed line represents GPCRs with *<* 50 known ligands, which accounts for 43.62% of total GPCRs.

To assess the performance of the one-shot learning architectures in predicting proteinligand binding, we apply three one-shot learning architectures to the kinase dataset. In image recognition, one-shot learning algorithms^22^ perform image classification tasks given only a few examples from each class. In binding ligand recognition, the similar learning task is to learn the behavior of new compounds given only a few data points from each kinase. In order to evaluate the performance with the different sizes of support set, we kept kinases with 50 positive inhibitors and 50 negative inhibitors. 183 unique kinases with 36, 628 inhibitors were retained in the input data set, and of these the first 80% kinase protein and remaining 20% were split as the training set (148 tasks) and testing set (35 tasks) respectively. AUC scores of the models on median held-out task are reported in Table 1. Three one-shot models obtained prediction performance of AUC score over 0.95.

**Table 1:** AUC score of Models on Median Held-out Task for Each Model on Kinase Collection.

| **Kinase** | **IterRefLSTM** | **AttnLSTM** | **Siamese** |
| --- | --- | --- | --- |
| 1+/1− | 0.980 ± 0.004 | 0.958 ± 0.075 | 0.943 ± 0.205 |
| 1+/5− | 0.963 ± 0.004 | 0.935 ± 0.214 | 0.958 ± 0.009 |
| 5+/10− | 0.989 ± 0.003 | 0.971 ± 0.010 | 0.981 ± 0.005 |
| 10+/10− | 0.989 ± 0.002 | 0.991 ± 0.004 | 0.986 ± 0.002 |
| 10+/20− | 0.991 ± 0.002 | 0.975 ± 0.004 | 0.992 ± 0.001 |
| 20+/20− | 0.986 ± 0.003 | 0.990 ± 0.004 | 0.986 ± 0.005 |
| 30+/30− | 0.993 ± 0.010 | 0.989 ± 0.005 | 0.993 ± 0.001 |

The reported AUC is mean ± standard deviation. The notation 1 + /1− means 1 positive sample and 1 negative sample in the support set. For each task, the evaluation was repeated 20 times, each time the support set is randomly selected.

Among three architectures, the iterative refinement LSTM displays a robust performance on kinase dataset. Furthermore, as the support size increases, the prediction performance also increased. Figure 3 shows the AUC score for each kinase protein in the testing set. For support set with 1+/1*−* (Figure 3a and 3b), IterRefLSTM had better prediction performance than AttnLSTM on 27/35 tasks. Similarly, IterRefLSTM demonstrated better performance than Siamese network on 22/35 tasks. For support set with 30 + /30*−* (Figure 3c and 3d), three architectures obtained similar AUC score on 30/35 tasks, but IterRefLSTM had very low AUC *<* 0.5 score on 4 tasks. In the predicted results with 30 + /30*−* support set, we found 4 outliers, IterRefLSTM had poor performance on these four kinases, and they are p38b (Mitogen-activated protein kinase p38b), p110*α* (phosphatidylinositol-4,5-bisphosphate 3-kinase, catalytic subunit *α*), p110*γ* (phosphatidylinositol-4,5-bisphosphate 3-kinase, catalytic subunit *γ*) and p110*δ* (phosphatidylinositol-4,5-bisphosphate 3-kinase, catalytic subunit *δ*), respectively. IterRefLSTM with 1 + /1*−* support set also showed weak predictive performance on p38b, p110*α* and p110*γ*. p110*α*, p110*γ* and p110*δ* are all from class I PI3Ks (phosphatidylinositol-4,5-bisphosphate 3-kinases) family, which are composed of catalytic subunits and regulatory subunits.

**Figure 3:**
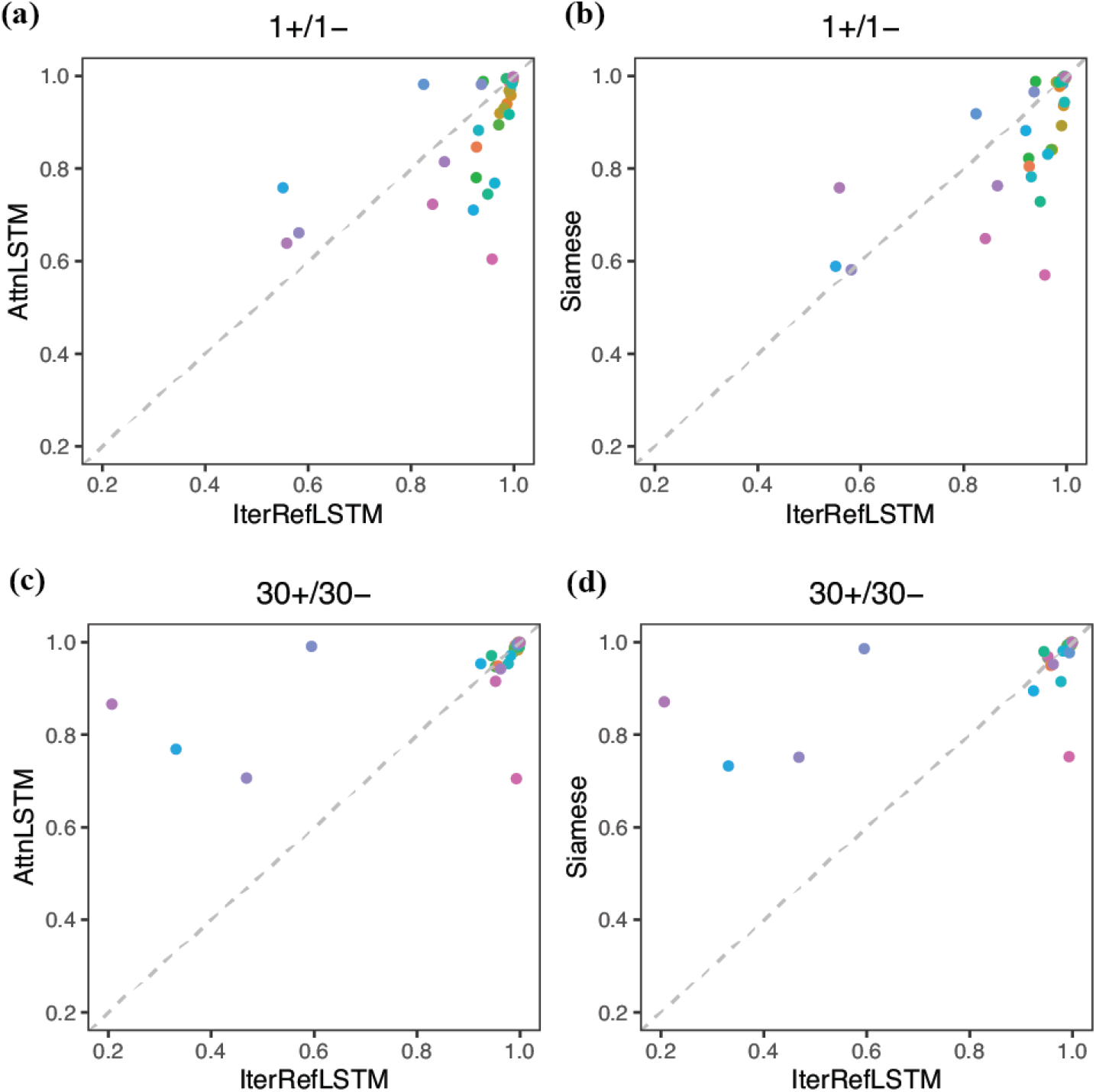
Comparison of AUC score of three one-shot models on 35 protein kinases. (*a*) and (*b*) are AUC score of three one-shot models with 1 + /1 *−* support set. (*c*) and (*d*) are AUC score of three one-shot models with 30 + /30 *−* support set. The different color dot represents one of 35 protein kinase in the test set.

### One Shot Learning Methods Provide Accurate Prediction of GPCR-Ligand Interaction

The GPCR dataset consists of 525 GPCRs with 132, 354 ligands, forming 215, 684 unique GPCR-ligand interactions. We observe that the majority of GPCR have a small number of known inhibitors (Figure 2b), among them, 43% of GPCRs have less than 50 known inhibitors and 20% only have less than 10 ligands.

To guarantee sufficient support set in the input data, we retained GPCRs with 50 positive inhibitors and 50 negative inhibitors. After this filtering step, 259 unique GPCRs with 132, 354 inhibitors were selected in the input data set. The data was split between test and training sets with 80% of GPCRs as the training set (208 tasks) and remaining 20% as the testing set (51 tasks). Table 2 shows AUC score of models on median held-out task for each model. Compared to the results from kinase dataset, we observe the prediction performance was lower with the highest AUC score is 0.799 from AttnLSTM model with 30 + /30*−* support set, whereas most AUC scores of prediction on kinase dataset were approximately 0.95 with smaller standard deviation. However, it is worth noting that an AUC score of 0.8 is considered an accurate prediction of GPCR-ligand interaction. The underlying reason of this drop in performance as compared to the kinases is likely the differences among the binding pockets of different sub-families of class A GPCRs. On the other hand, kinases have a highly conserved ATP binding pocket, which is targeted by the inhibitors. In addition, the input data for GPCR, not only includes GPCRs in human, but also GPCRs from other organisms, like rat, pig and rabbit etc. Therefore, transferability of knowledge across different types of GPCRs is expected to be limited.

**Table 2:** AUC score of Models on Median Held-out Task for Each Model on GPCR Collection.

| GPCR | IterRefLSTM | AttnLSTM | Siamese |
| --- | --- | --- | --- |
| 1+/1− | 0.624 ± 0.092 | 0.695 ± 0.199 | 0.624 ± 0.092 |
| 1+/5− | 0.584 ± 0.085 | 0.773 ± 0.091 | 0.584 ± 0.085 |
| 5+/10− | 0.620 ± 0.143 | 0.780 ± 0.069 | 0.620 ± 0.144 |
| 10+/10− | 0.649 ± 0.044 | 0.730 ± 0.084 | 0.649 ± 0.044 |
| 10+/20− | 0.634 ± 0.056 | 0.740 ± 0.061 | 0.634 ± 0.056 |
| 20+/20− | 0.640 ± 0.090 | 0.747 ± 0.032 | 0.640 ± 0.090 |
| 30+/30− | 0.638 ± 0.010 | 0.799 ± 0.046 | 0.638 ± 0.010 |

The reported AUC is mean ± standard deviation. The notation 1 + /1− means 1 positive sample and 1 negative sample in the support set. For each task, the evaluation was repeated 20 times, each time the support set is randomly selected.

For the performance on the GPCR collection, the AttnLSTM has a better prediction results in most tasks and increased size of support set improves the AUC score. Figure 4 shows the AUC score for each GPCR in the testing set. For support set with 1+ /1*−* (Figure 4a and 4b), AttnLSTM has a better prediction performance than IterRefLSTM on 41/51 tasks, and the Siamese network on 47/51 tasks. For support set with 30 + /30*−* (Figure 4c and 4d), AttnLSTM has higher AUC score than IterRefLSTM on 42/51 tasks, and than Siamese network on 47/51 tasks.

**Figure 4:**
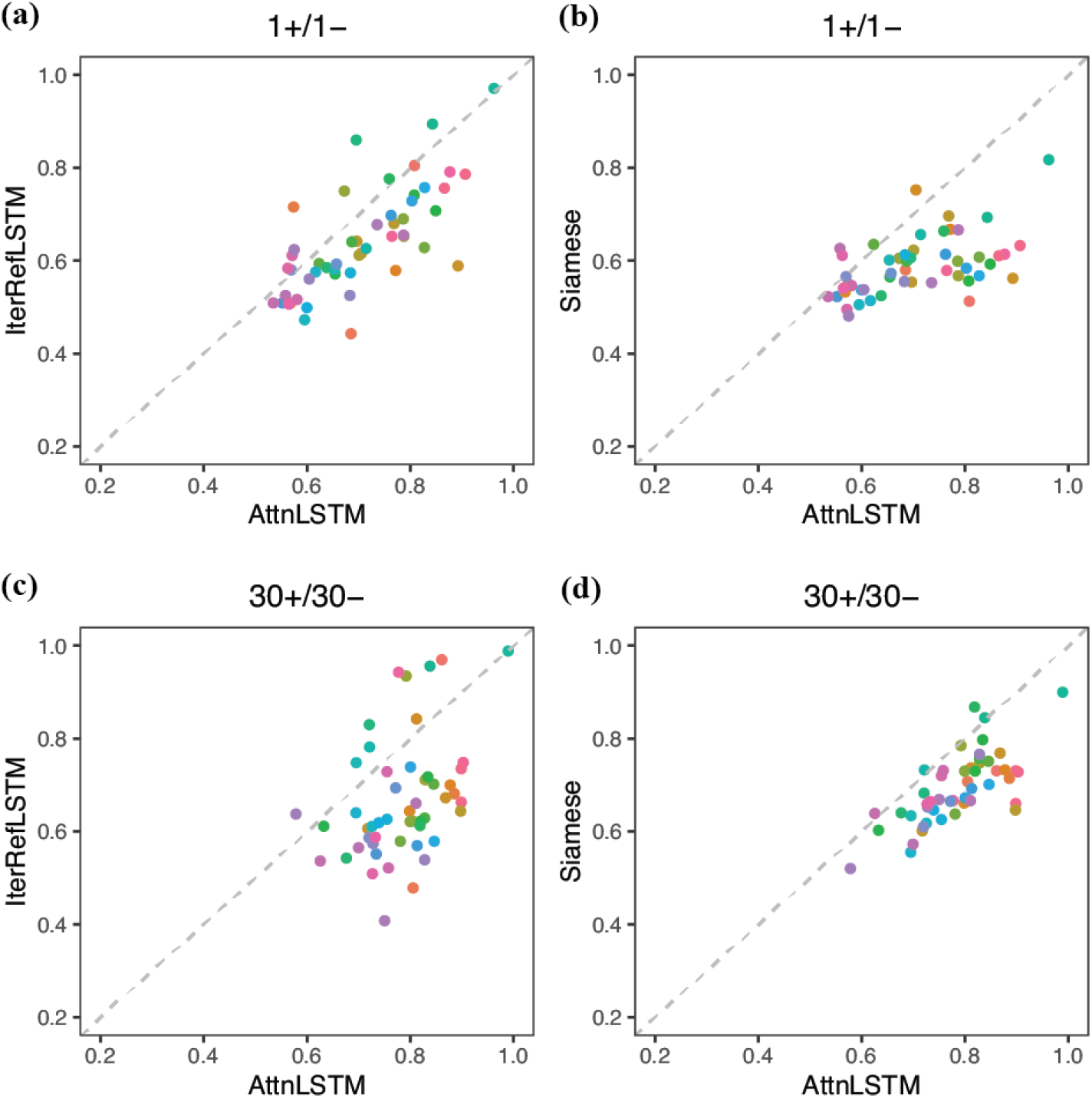
Comparison of AUC score of three one-shot models on 51 GPCRs. (*a*) and (*b*) are AUC score of three one-shot models with 1 + /1 *−* support set. (*c*) and (*d*) are AUC score of three one-shot models with 30 + /30 *−* support set. The different color dot represents one of 51 GPCRs in the test set.

### Predicting the biological activities of excipients

On the basis of the performance of different one-shot learning architectures on kinases, IterRefLSTM was found to provide the highest AUC score. Therefore, we train a IterRefLSTM using 30 + /30*−* support set on the entire Kinase collection and apply the model to predict the interaction between kinases and excipients. The list of excipients were retrieved from Reker’s paper,^10^ 799 unique excipients were included as prediction tasks. All predicted binding probability of kinase and each excipient were shown in the Table S1. Figure 5a shows a list of 50 excipients which were predicted to bind to the most kinases.

**Figure 5:**
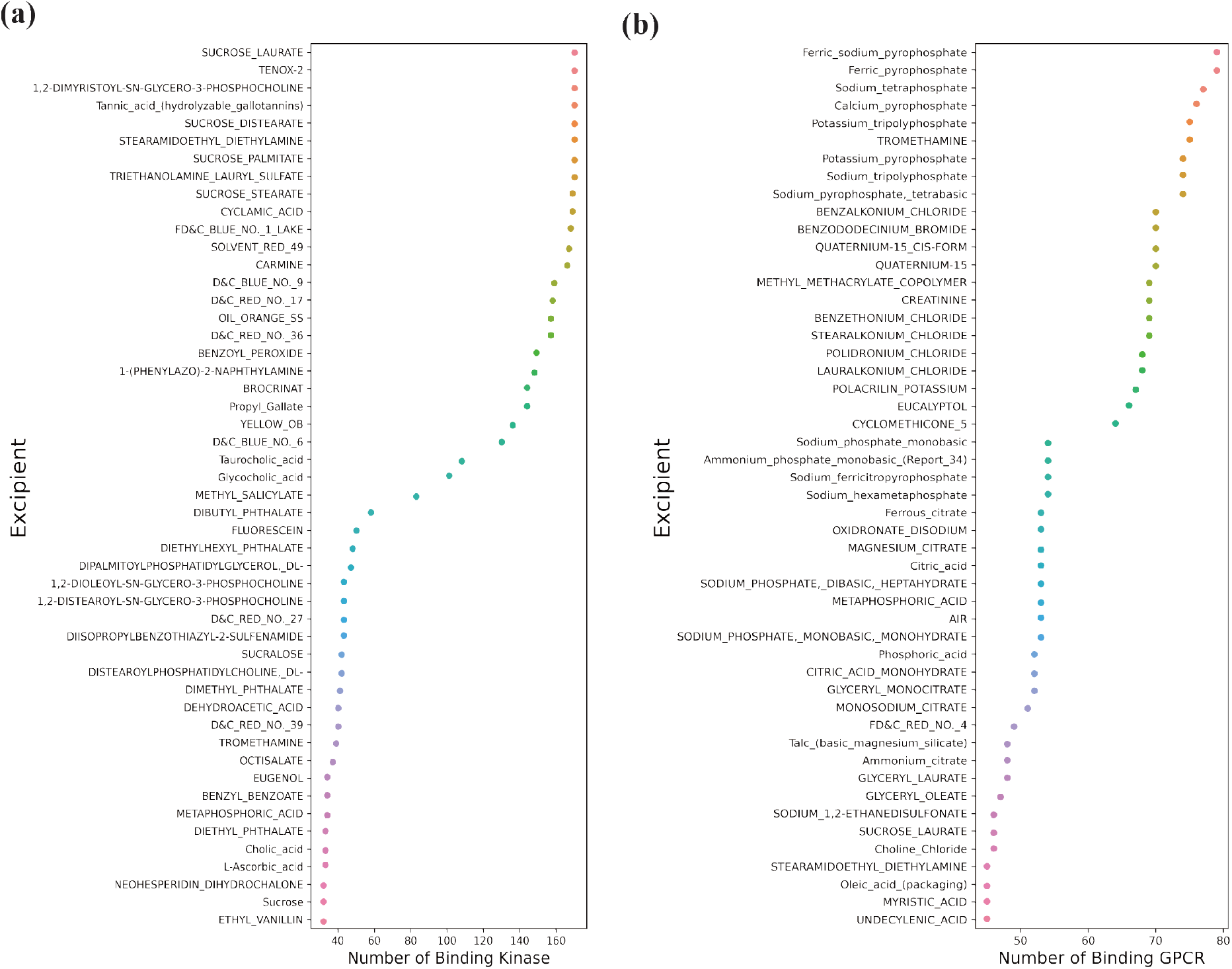
Top 50 excipients binding to the most kinases/GPCRs.

Tannic acid is a kind of hydrolysable tannins that is widely exists in tea, fruits, vegetables, red wine and coffee.^30^ As an important part of the human diet, investigating the biological effects of tannic acid is significant meaningful for formulations design of food/drug. The known targets of tannic acid include MAP kinase Erk1, MAP kinase Erk2, Tyrosine-protein kinase FYN, Tyrosine-protein kinase LCK, MAP kinase p38 alpha.^10^ For these targets, the predicted probability of interaction is 0.613, 0.706, 0.757, 0.710 and 0.793, respectively. The predicted results from our trained one-shot model keep consistent with experimental results. On the other hand, we also investigated for a specific kinase whether predicted binding compounds would have similar structure with the known inhibitor. Epidermal growth factor receptor (EGFR) is an key receptor tyrosine kinase which has been shown to prevent tumor cell death by maintaining the basal intracellular glucose level.^31^ Erlotinib (TARCEVA) is a FDA-approved reversible inhibitor of EGFR tyrosine kinase that functions as an ATP analogues by competing with ATP binding pockets and thereby inhibiting further cellular proliferation, blocking cell-cycle.^32^ Figure 6a shows the structure of EGFR complexed with erlotinib (PDB code: 1M17^33^) and Figure 6b display the 2D structure of erlotinib. Among the excipients predicted to bind to EGFR, three representative compounds are shown in Figure 6c, including D&C Red No.17, Oil Orange SS and D&C Red No.36, all used as dyes for coloring medicines so that they can be better distinguished by patients. These three excipients have similar chemical groups and display similar structure with known EGFR inhibitor erlotinib, which give us more confidence for the prediction results from one-shot models.

**Figure 6:**
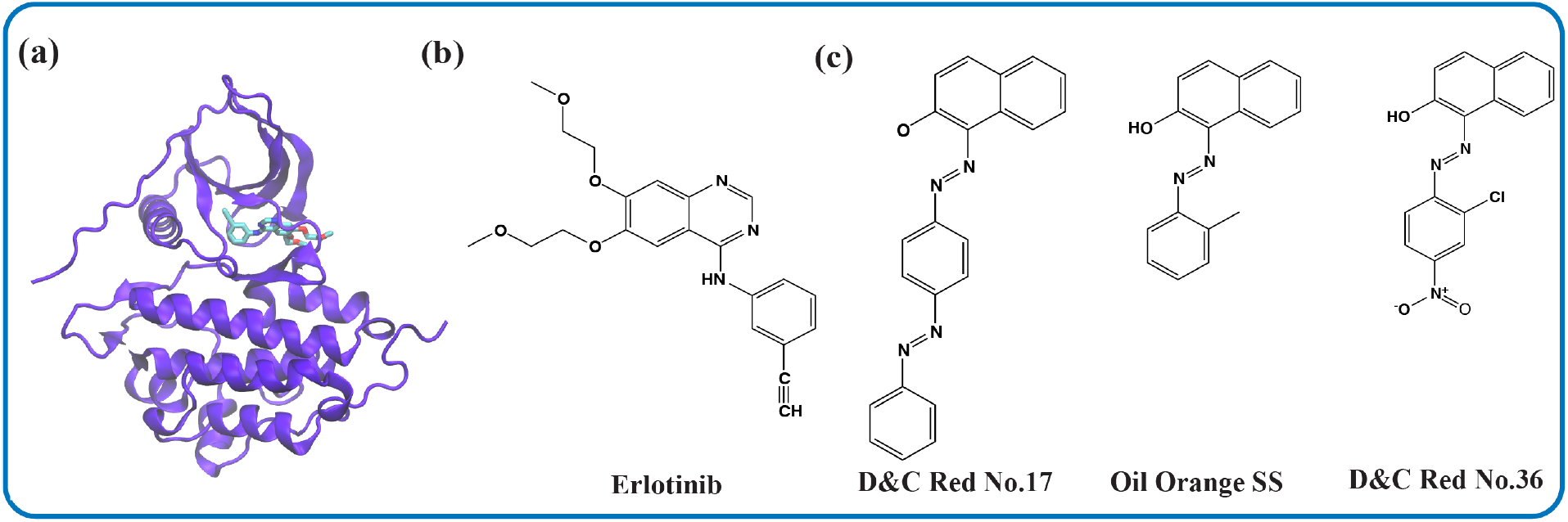
Predicted excipients binding to EGFR. (*a*) The structure of EGFR shown in purple cartoon representation and Erlotinib in blue sticks (PDB code: 1M17^33^). (*b*) The chemical structure of known EGFR inhibitor, Erlotinib. (*c*) Chemical structures of representative excipients binding to EGFR from prediction

For GPCR collection, AttnLSTM had the best performance as compared to the other two one-shot learning architectures. Therefore, we train a AttnLSTM using 30 + /30*−* support set on the entire GPCR collection, and apply the model to predict the possible excipients binding to GPCRs. All predicted binding probability of GPCR and each excipient were shown in the Table S2. Figure 5b shows a list of 50 excipients which were predicted to bind to the most GPCRs.

Histamine H1 receptor (HRH1) is a typical rhodopsin-like GPCR which can be activated by biogenic amine histamine. By activating HRH1, histamine is involved in many pathophysiological processes, like allergies and inflammations.^34^ As a result HRH1 antagonists are proved as effective drugs alleviating allergic reactions. Doxepin is approved drug as effective HRH1 antagonist. Figure 7a shows the structure of HRH1 complexed with doxepin (PDB code: 3RZE^34^) and Figure 7b display the chemical structure of doxepin. In Figure 7c, we show chemical structures of three compounds which were predicted with highest binding probability with HRH1. These three compounds, Lauralkonium chloride, Stearalkonium chloride and Benzalkonium chloride have similar structure and they are safe used as cosmetic ingredients. In addition, we also compared predicted results of GPCR and excipients/GRAS with known biological activities. Benzethonium chloride can function as antimicrobial preservative, antiseptic and disinfectant, its known targets are Dopamine receptor D1 and Histamine Receptor H3.^1^ The predicted binding probability for these two targets is 0.633 and 0.579, respectively, which are both consistent with experimental results.

**Figure 7:**
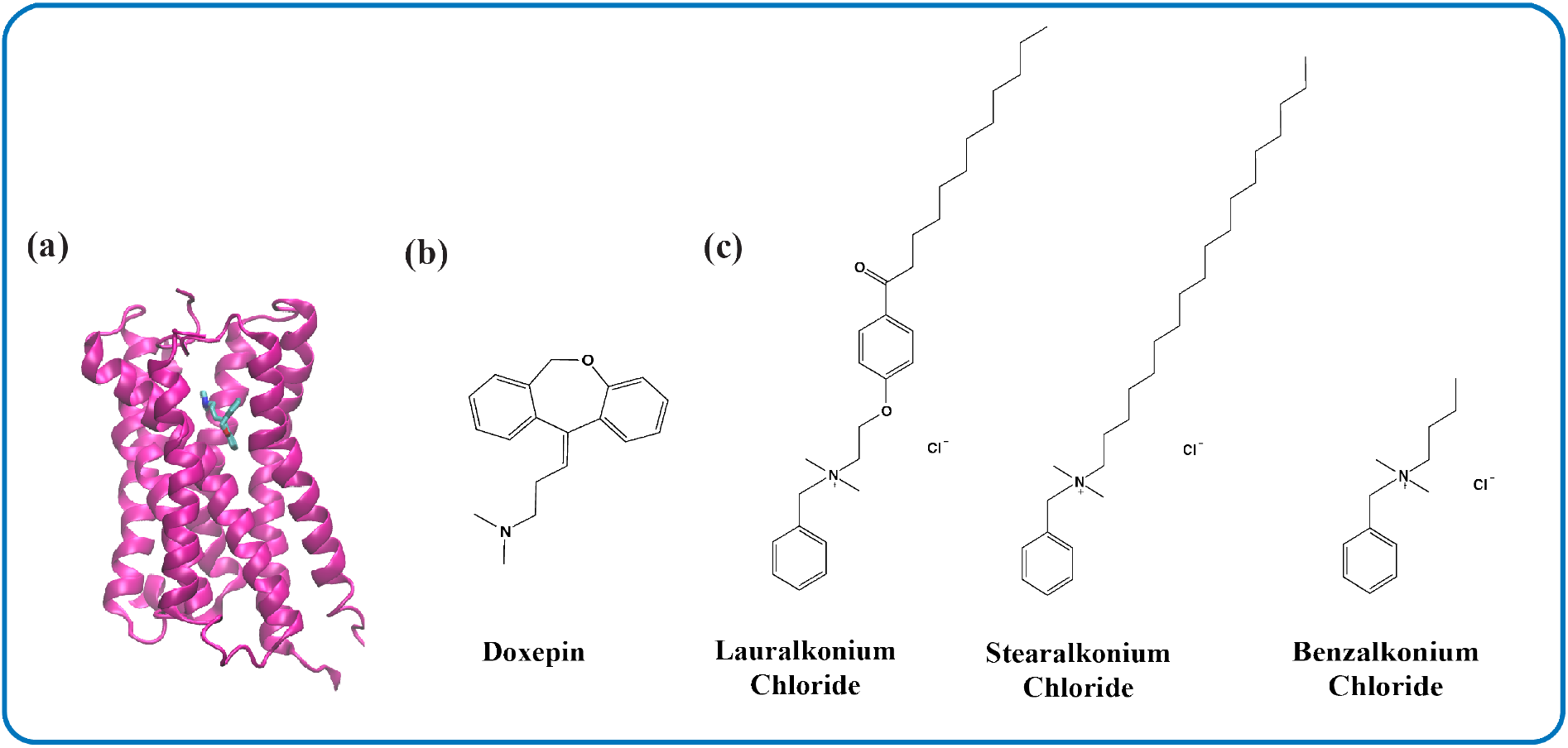
Predicted excipients binding to HRH1. (*a*) The structure of HRH1 shown in magenta cartoon representation and Doxepin in blue sticks (PDB code: 3RZE^34^). (*b*) The chemical structure of known ligand of HRH1, Doxepin. (*c*) Chemical structures of representative excipients binding to HRH1 from prediction.

## Discussion

In this work, one-shot learning models are first applied to the kinase collection and GPCR collection. Due to limited known compounds binding to protein kinase and GPCRs, traditional deep learning models are not widely applied in these dataset. For kinase collection, three one-shot models shows accurate prediction performance, while for GPCR dataset, the prediction achieve a little bit lower accurate predictions. The possible underlying reason is that the ligand binding domains in class A GPCRs are various. Figure 8 shows the diversity of ligand binding domain in class A GPCR, such as nucleoside binding domain, acetylcholine binding domain, lipids binding domain, peptide binding domain.^35^ Alternatively, the ATP binding pocket in kinase is highly conserved.^36^ For example, in Figure 9, the WebLogo plot was generated to show the binding pocket sequence conservation for 36 Kinase pockets in CAMK (cAMP-dependent protein kinase) group.^37^ The overall height of the sequences in WebLogo represents the frequency and conservation at the corresponding position. Based on the WebLogo, the G2, V4, K6, E7, N24, F29 residues are highly conserved. Therefore, there is less structural similarity between ligands for each GPCR compared to protein kinase. One-shot learning models require some extent structural similarity in order to extrapolate from limited data, and classify new compounds.

**Figure 8:**
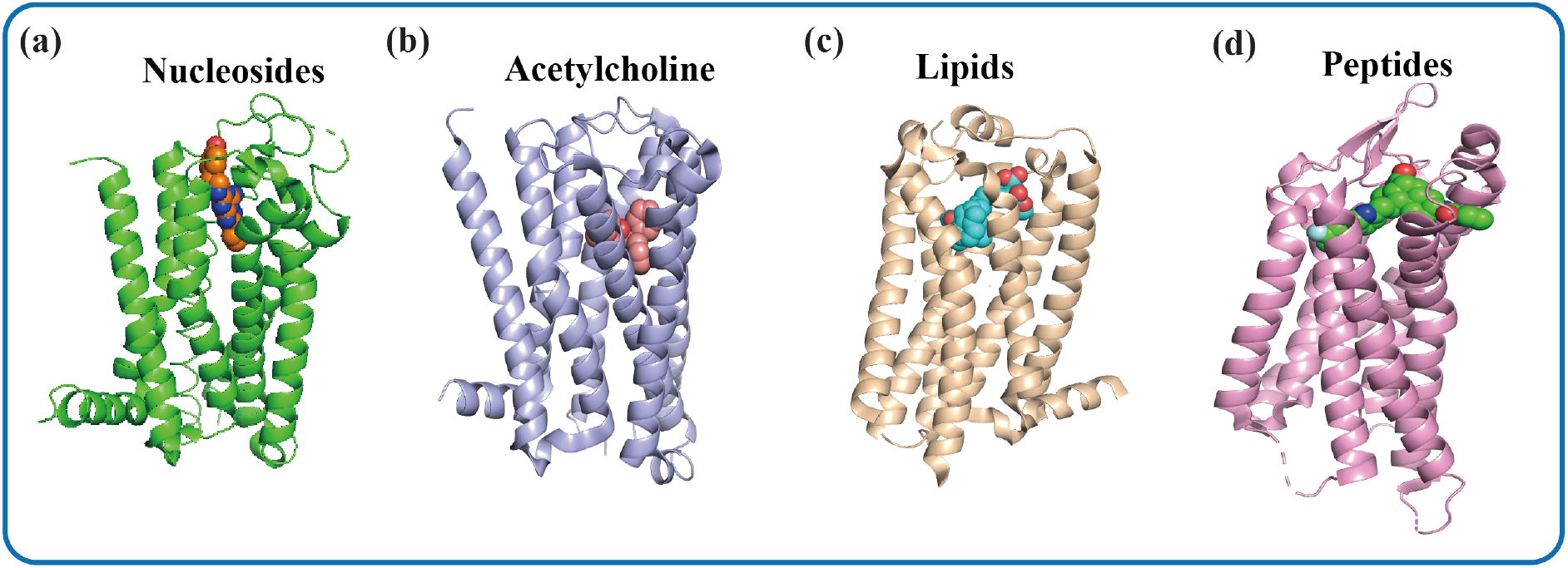
Diversity of ligand binding domain in class A GPCR. (*a*) The structure of human A2A adenosine receptor bound to ZM241385 (PDB code: 3EML^38^). (*b*) The structure of human M2 muscarinic acetylcholine receptor bound to QNB (PDB code: 3UON^39^). (c) The structure of human Lysophosphatidic Acid Receptor 1 bound to ON9 (PDB code: 4Z35^40^). (*d*) The structure of human protease-activated receptor 1 (PAR1) bound to VPX (PDB code: 3VW7^41^).

**Figure 9:**
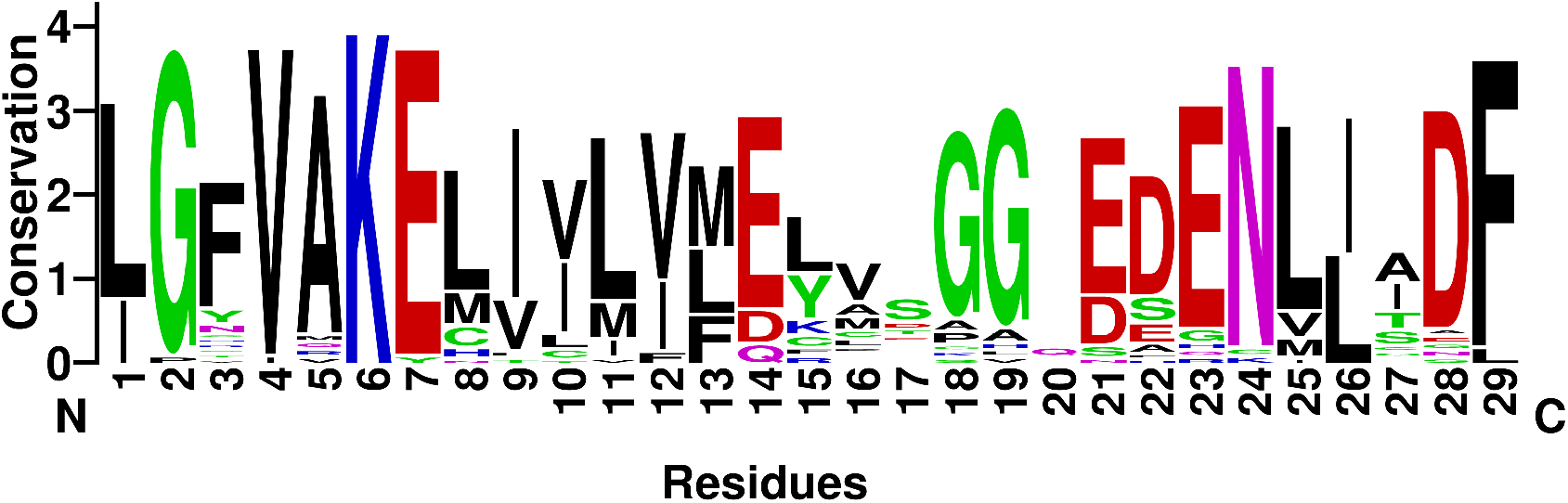
The sequence WebLogo of 36 Kinase binding pockets in CAMK (cAMP-dependent protein kinase) group.

Another aspect of our work is predicting the interaction between excipients and protein kinase or GPCRs. Excipients are necessary as support to the active ingredients in drug and play key role in drug’s stability, preservation, appearance, pharmacokinetics and bioavilability.^42^ However, excipients can trigger adverse events in patients. A small number of patients are allergic to drug excipients, food derivatives or dyes.^2,43,44^ Excipients containing gluten may result in adverse effects in the patients with celiac disease or gluten sensitivity.^45^ Therefore, a large number of different versions of the same drug with different excipients are currently available. The comprehensive investigation of biological activities of excipients is significant meaningful to excipients design. Our work provides an efficient strategy for identifying possible binding excipients to two most important collections of drug targets.

Considering the biological effects of active and inactive ingredients may be helpful to drug formulation design and refine clinical decision to control adverse effects and the pharmacokinetics of drugs.

Overall, we anticipate that one-shot learning models will be useful in predicting the interaction of ligands to receptors with limited available data, especially for receptors which have more conserved binding pocket. Typically, excipient formulations are designed to improve the drug stability, bioavailability and other metrics.^46–50^ Here, our work suggests a deep learning architecture used for identifying biological targets of drug excipients which could help to design the most suitable excipients to provide personalized medicine to a patient and avoid possible adverse effects from inactive ingredients.

